# An Information Geometry approach to model topological trajectories and Gene Expression Radius from UMAP geometry

**DOI:** 10.64898/2026.08.27.747659

**Authors:** Marco Polo Castillo-Villalba, Froylán Espinosa Bustamante

## Abstract

Understanding the relationship between gene expression dynamics and cellular identity remains a central challenge in single-cell biology. Here, we introduce a novel computational and mathematical framework that integrates information geometry, fuzzy topology, and UMAP analysis to model gene expression landscapes derived from single-cell RNA sequencing data.

We formalize gene expression data as a fuzzy topological space, where interactions between expression points are governed by probabilistic distributions inspired by manifold learning approaches such as UMAP. Within this framework, we define an information geometric structure through a Fisher metric induced by these distributions, enabling the computation of geodesic trajectories that capture cellular differentiation processes.

A key contribution of this work is the derivation of analytical conditions, expressed as expression radius formulas, that characterize local neighborhoods in gene expression space. These conditions allow for the identification of genes associated with stem cell states and predictions in transitional cell types in future work.

Application of the proposed framework to single-cell datasets reveals biologically meaningful gene sets enriched in key regulatory pathways and transcription factors, demonstrating the capacity of our approach to uncover latent structure in complex gene expression data.

Our results suggest that integrating differential geometry with statistical learning theory offers a powerful paradigm for modeling genotype–phenotype relationships and cellular state transitions, with potential implications for precision medicine and systems biology.

## 1. Methods: Information Geometry in UMAP and Fuzzy Topology

In this project, we characterize differential expression points derived from single-cell RNA sequencing (scRNA-seq) data as a one-dimensional fuzzy line, also referred to as a fuzzy 1-simplex. That is, given two expression points *Y*_*i*_ and *Y*_*j*_ in the logarithmic space log_10_, which quantifies changes in gene expression (fold change), the probability of interaction between these points is weighted by the distributions *V*_*ij*_ for differentiated cell types and *W*_*ij*_ for stem or progenitor cell types.

A fuzzy line, or fuzzy 1-simplex, between clusters of differential expression points *Y*_*i*_ and *Y*_*j*_ can be quantified by the **expression radius**, or fractional distance, defined as:

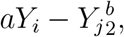

where *a* and *b* are model hyperparameters, which are related to the standard deviation of expression points. This metric induces and formally defines a fuzzy topology that characterizes the entire expression data space. The open sets or neighborhoods are defined by these radii for each pair of points *Y*_*i*_ and *Y*_*j*_.

### 1.1. UMAP (Background): Uniform Manifold Approximation and Projection

In this project, for the analysis of expression data derived from single-cell sequencing, we have extensively used the UMAP algorithm, see [2], and PAGA, see [3]. From these methods, we adopt the validity of the probability distributions that characterize different cell types, as described below:

**Conditional distribution for the interaction between two expression points** *Y*_*i*_ **and** *Y*_*j*_ **in differentiated cell types:**

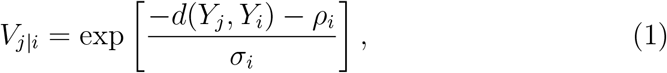

**Joint distribution for two expression points** *Y*_*i*_ **and** *Y*_*j*_ **in differentiated cell types:**

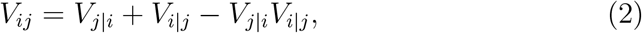

**Distribution characterizing the interaction between two gene expression points** *Y*_*i*_ **and** *Y*_*j*_ **in stem or progenitor cells:**

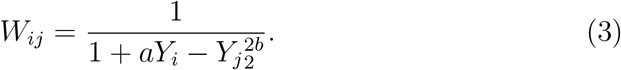

From these probability distributions, we construct information-geometric structures and define geodesics or trajectories of points on the fuzzy topology that characterize gene expression data derived from single-cell techniques.

### 1.2. Information Manifolds or Statistical Learning Manifolds

A differentiable manifold consists of a topological space defined by a collection of open sets (local neighborhoods), such that finite unions and pairwise intersections of these sets also belong to the topology. There exist bijective mappings between these open sets that are continuous and infinitely differentiable. The composition of these mappings and their inverses defines coordinate transformations on the intersections of these neighborhoods.

When a multilinear mapping exists between coordinate points in the tangent spaces of these neighborhoods and the real numbers, such a mapping is called a tensor. When a tensor defines a metric that measures distances between points, the structure described above defines a Riemannian manifold. In the case of information manifolds, the points of the manifold are characterized by probability distributions, as in the case of gene expression data from single-cell experiments. A distance between distributions of points is then defined, for instance, between those described by *V*_*ij*_ and *W*_*ij*_ across different clusters. This distance is given by the information entropy, or Kullback–Leibler divergence, as shown below.

**Information Entropy and Fisher Information Matrix**

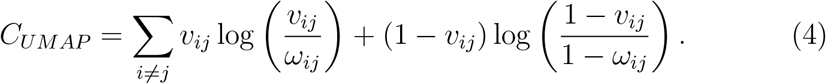

This probabilistic distance between clusters induces the construction of a metric tensor, known as the Fisher information matrix. Furthermore, this object defines a Riemannian manifold, commonly referred to in the literature as an information manifold.

The concept of an information or statistical learning manifold arises from the probability distributions *V* and *W* that characterize gene expression data through UMAP.

**Fisher Matrix and Distance Between Expression Clusters**

The line element that separates two points in Euclidean space is given by:

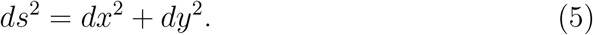

In the case of a fuzzy topology and an information manifold, the line element (distance) between two distributions that separate clusters of data—both for stem cells and differentiated cell types along a fuzzy trajectory—is given by:

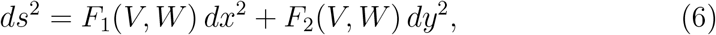

where the coefficients *F*_1_(*V, W*) and *F*_2_(*V, W*) are stochastic functions depending on the distributions *V* and *W*. These correspond to the diagonal elements of the Fisher information matrix, namely *I*_11_ and *I*_22_.

**Infinitesimal Representation of the Fisher Information Matrix**

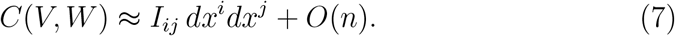

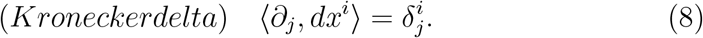

By applying the contravariant differential operator and using the orthogonality property of the Kronecker delta, we obtain:

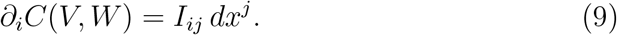

Applying the differential operator once again with respect to the index *j*, we obtain an explicit expression for computing the Fisher information matrix in the infinitesimal case:

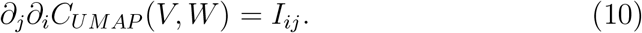

### 1.3. Computation of Fuzzy Trajectories, Geodesics and Expression Ratios

Using the geodesic equation on a Riemannian information manifold, we obtain the following differential equation for the computation of geodesics:

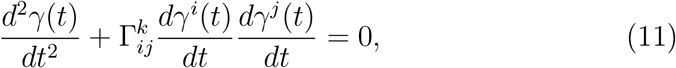

where the connection functions 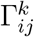 are the Christoffel symbols, which

describe the curvature and parallel transport of information vectors along a trajectory. The function *γ*(*t*) analytically represents a trajectory on the information manifold.

In the case of the symmetric Fisher information tensor, when it is diagonal for single-cell expression data, the global trajectory is governed by the following differential equation:

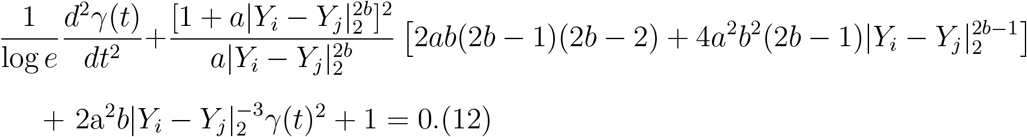

At present, an analytical solution for these trajectories is not tractable. However, by exploiting the local fuzzy topology, we focus on solutions defined over a fuzzy 1-simplex, which provide a local approximation of trajectories. In this case, the solutions reduce to fuzzy straight lines, given by:

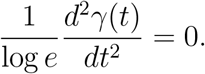

A relevant condition for such geodesic solutions is that two infinitesimally close differential expression points *Y*_*i*_ and *Y*_*j*_ have a line element satisfying *ds*^2^ ≈0 (see Eq. A.2). This implies that the coefficients of the Fisher information matrix vanish:

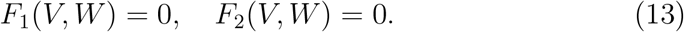

It is important to note that the expression for *F*_1_ involves exponential functions, which do not vanish unless the variance of the expression data becomes very large, as is the case for highly differentiated cell types. Therefore, the most informative condition arises from *F*_2_(*V, W*) = 0.

Assuming that two differential gene expression points lie infinitesimally close within a fuzzy 1-simplex neighborhood, we obtain the condition:

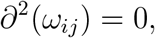

which leads to the **expression radius formula for stem cells**. This formula is derived from the *I*_22_ entry of the Fisher information matrix:

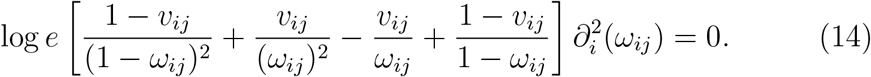

Since the probability terms in brackets cannot vanish simultaneously, we obtain:

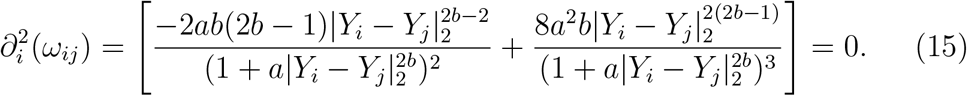

Solving for the differential expression radius |*Y*_*i*_ −*Y*_*j*_ | _2_, we obtain a Expression Ratio:

**Gene Expression Radius**

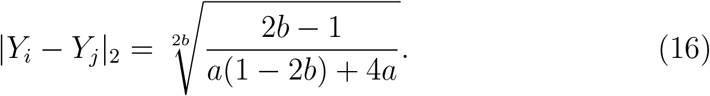

Substituting the standard UMAP parameters *a* = 1.929 and *b* = 0.7915, we obtain the numerical radius for fuzzy neighborhoods associated with stem-like cell types:

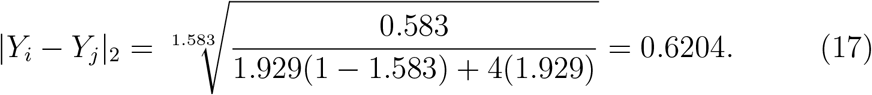

That is, all gene expression data satisfying the condition |*Y*_*i*_ − *Y*_*j*_ | _2_ = 0.6204 correspond to pairs of points that are infinitesimally close and connected by a fuzzy 1-simplex. When characterized by the source distribution *w*_*ij*_, these points are associated with stem or progenitor cell types and represent the initial genes expressed along a geodesic trajectory. These trajectories can be interpreted as velocity flows of transcripts, since fuzzy 1-simplex elements correspond to solutions of the geodesic equation on the Riemannian information manifold.

## 2. Formulas that characterize transitions between cell types

Returning to the estimation of the expression radius formulas, we base our analysis on the following hypotheses, which provide an initial bound for the first set of genes associated with changes in differential expression radii.

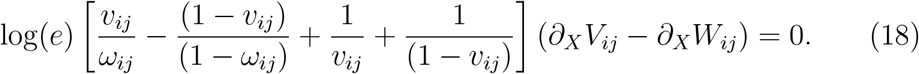

In the context of theoretical physics, differences in gradients can be associated with boundary conditions of the phenomenon under study, such as differences in electric potential or pressure across a given geometry (e.g., cylindrical domains). In this work, we are particularly interested in the cell cycle of organoids *in vivo*, where the correct interpretation of these gradient differences plays a key role.

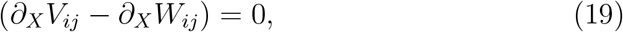

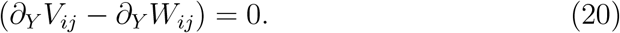

Quantitative proteome analysis leads to the continuity equation:

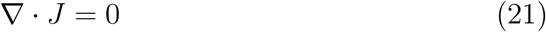

**General case for transitions between cell types**.

In this case, we consider:

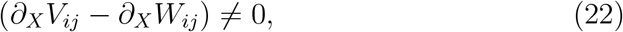

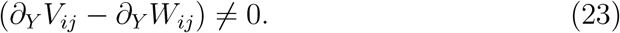

From this, we obtain the following set of expressions:

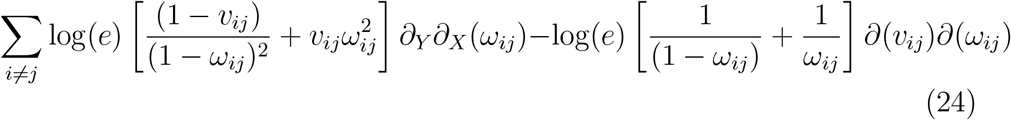

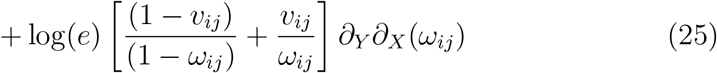

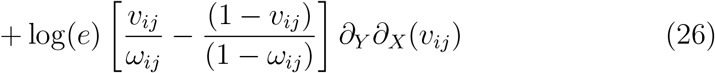

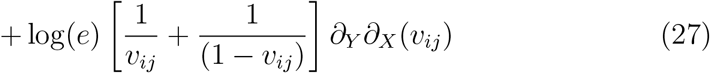

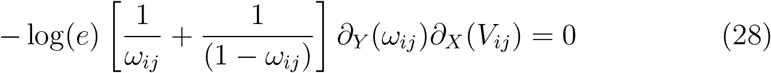

**Hypothesis: zero flux conditions during transitions between cell types**.

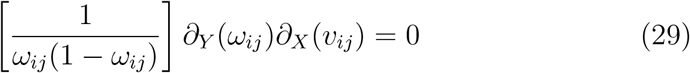

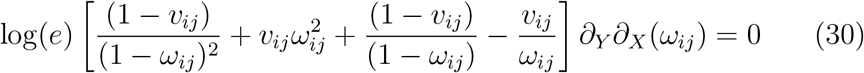

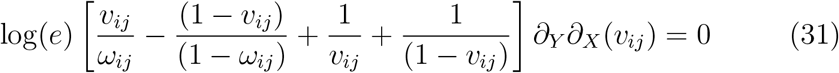

**By summing the previous expressions, we obtain:**

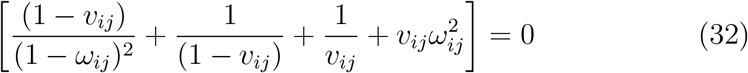

Solving for the distribution *v*_*ij*_, we obtain:

**General expression for cell-type transitions between stem and differentiated states: Implies different expression ratios**.

**Expression Radius for Transition Cell-Types**.

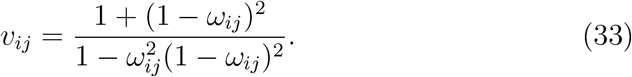

This expression represents a mixture of probability distributions corresponding to both cellular states (stem and differentiated). Next, we perform a Taylor series expansion. By expanding up to fourth order and matching coefficients, we obtain new values for the hyperparameter *b* and expressions for the hyperparameter *a*.

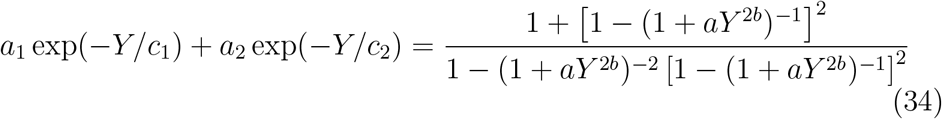

Rewriting:

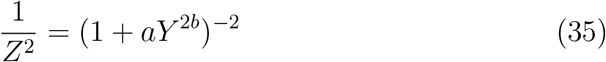

Further simplification yields:

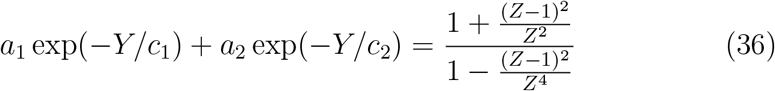

**Series expansion:**

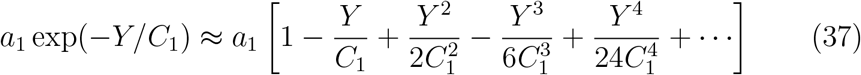

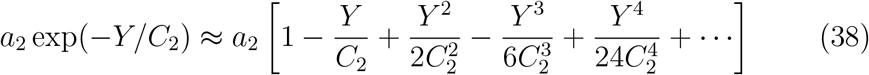

**We obtain different cases of Expression Radius depending on the value of** *b*:

**Case** *b* = 1:

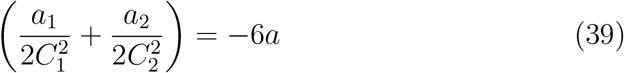

**Case** *b* = 3*/*2:

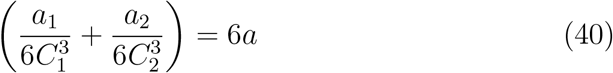

**Case** *b* = 1*/*2:

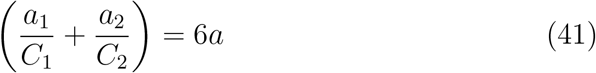

**Case** *b* = 2:

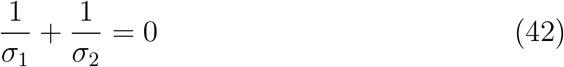

## 3. Results

### 3.1. Metabolic pathway enrichment for genes satisfying the expression radius condition F_2_(V, W) = 0 and equation (16) for stem cells

A total of 530 genes were obtained, of which 391 showed significant enrichment in relevant metabolic pathways using the Enrichr platform. The analysis identified global transcription factors such as **USF2, MYC, JUND**, and **MAZ**, with statistically significant p-values of 0.001314, 0.007637, 0.01055, and 0.01253, respectively. An example of the enrichment analysis for the transcription factor USF2 is presented below.

### 3.2. Conclusions

In this work, we have presented an analytical framework based on the potential differential geometry underlying the geometric structure of UMAP. The subsequent analysis opens a new framework for computing trajectories and geodesics associated with different cell-type states in scRNA-seq datasets. At the same time, our framework provides a new set of mathematical formulations for characterizing families of genes within fuzzy neighborhoods, that we named as: **Gene Expression Radius**. These groups of genes may provide insights into the identification and characterization of distinct cell types. A more in-depth analysis is required before this work can be submitted to a peer-reviewed journal.

## Supplementary files

-We provide the list of genes satisfying the expression radius condition given by equation (16) in an Excel file named: *Gene*_*E*_*xpression*_*R*_*adius*_*A*_*nalysis.xlsx*.

*-For the computation of expression radii using the formula* |*Y*_*i*_ − *Y*_*j*_ | _2_ = 0.6204, *the full set of gene combinations is reported in the file Combinatorial-genes*_*e*_*xpression*_*r*_*adii.txt*.

---

**Figure 1:**
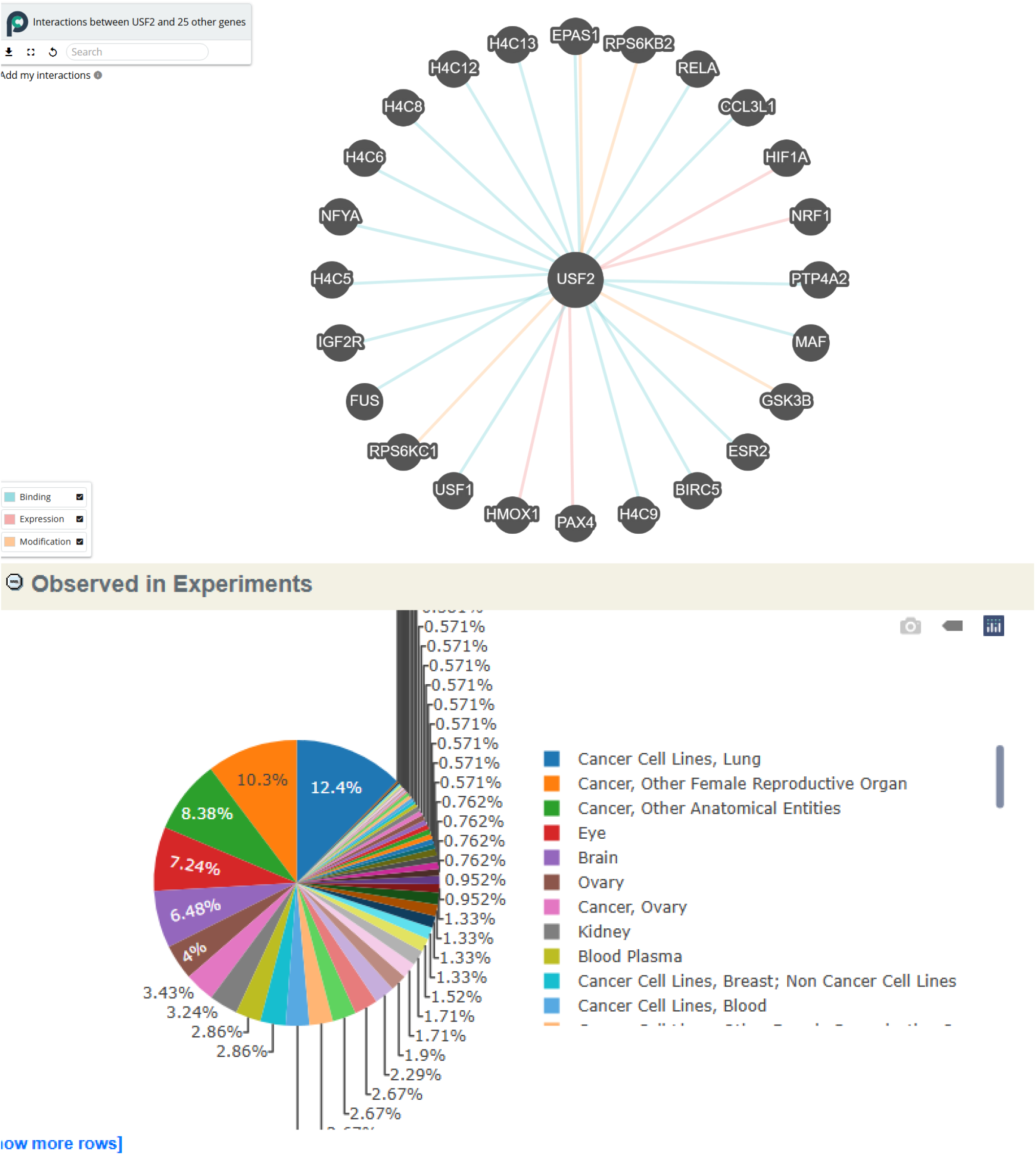
Metabolic pathway enrichment for 391 genes obtained from the expression radius defined in (16) for stem cells.

